# Giant Vesicle Platform for Synthetic Endoplasmic Reticulum Networks

**DOI:** 10.64898/2026.10.10.756113

**Authors:** Cade Vallero, Shreya Pramanik, Tsu-Wang Sun, Miroslaw Tarnawski, Joachim P. Spatz, Rumiana Dimova, Reinhard Lipowsky

**Author notes:** Correspondence should be addressed to R.L.

## Abstract

The endoplasmic reticulum (ER) is a membrane-bound organelle found in all eukaryotic cells. This organelle has a fascinating morphology, which involves a network of membrane nanotubes and three-way junctions connecting three nanotubes. Previous reconstitution studies were based on proteo-liposomes with a size of 100 to 200 nanometers, using electron microscopy as the main method to image the resulting membrane morphologies. These studies revealed the importance of membrane GTPases, that is, membrane proteins that hydrolyse GTP. Here, we introduce an alternative platform for the formation of synthetic ER networks starting from giant unilamellar vesicles with a typical size of 10 mirometers and containing a single membrane GTPase such as atlastin from fruit flies or Sey1 from yeast. This giant vesicle approach makes the networks accessible to optical microscopy, by which one can elucidate both their structure and their dynamics. We first demonstrate that the membranes’ propensity to form synthetic ER networks depends strongly on the lipid composition. More precisely, we show that this propensity varies non-monotonically with the mole fraction of cholesterol and that the frequency of three-way junctions has a maximum for an optimal mole fraction between 10 and 15 mol%. This optimal value is comparable to the mole fraction of cholesterol as estimated for ER membranes in vivo. We also demonstrate the formation of synthetic ER networks when the membranes contain only the membrane GTPase Sey1 but no other membrane protein, which implies that Sey1 is able on its own to generate a high membrane curvature. Based on the reconstitution approach introduced here, the synthetic ER networks can now be further investigated using the versatile toolbox that has been developed for giant vesicles.

## 1 Introduction

The endoplasmic reticulum (ER) is the largest membrane-bound organelle, found in all eukaryotic cells. Each cell contains only a single copy of this organelle, which represents, however, more than half of the total cellular membrane [1]. The morphology of the ER membrane involves several subregions: the perinuclear subregion close to the nucleus consists of many sheets whereas it forms narrow nanotubes in the peripheral subregion further away from the nucleus. [2] The membrane nanotubes are interconnected by three-way junctions [3–6] and form a continuous three-dimensional network that extends throughout the whole cell. [2, 7] During mitosis, the ER membrane is significantly remodeled: it moves out to the cell periphery and its morphology becomes dominated by nanotubes. [8–11]

In vivo, the ER nanotubes have a typical diameter between 50 and 100 nm. [2, 12] The high curvature of the nanotubes is thought to be generated by two protein families, the reticulons and the receptor expression enhancing proteins (REEPs) [13, 14] In the context of curvature elasticity, the formation of nanotubes implies a large spontaneous curvature, which determines the diameter of the tubes. [15, 16] The complex nanotubular network of the ER is formed by a single membrane, enclosing a continuous nanofluidic network of water channels, as demonstrated by monitoring the diffusion of fluorescently labeled molecules. [2, 17, 18] As a consequence, the ER membrane represents a bicontinuous structure that partitions the intracellular space into two separate, interpenetrating subcompartments, forming a surface with a high topological genus. [19]

In-vitro reconstitution experiments based on cell extracts [20] and proteo-liposomes [21–25] have demonstrated that synthetic ER networks can be formed in the absence of any cytoskeletal elements. On the other hand, in order to form such networks, the proteo-liposomes must contain a membrane GTPase, that is, a membrane protein that is able to hydrolyze GTP. [21–23,25–28] In vivo, ER networks with three-way junctions are formed by membrane fusion [5], which is coupled to GTP hydrolysis and mediated by the trans-dimerization of a membrane GTPase. So far, synthetic ER networks have been obtained using proteo-liposomes with the membrane GTPase atlastin from *Drosophila melanogaster* [21–23, 29] or with two proteins from the yeast *Saccharomyces cerevisiae* as provided by the membrane GTPase Sey1 and the membrane protein Yop1 that generates membrane curvature [22, 23, 25, 28]. The membrane morphologies formed by the proteo-liposomes have been imaged by variants of electron microscopy. All electron microscopy methods are, however, restricted to a single snapshot of each proteo-liposome and cannot monitor the time-dependent behavior and morphological transformations of individual liposomes.

Here, we introduce and develop an alternative platform for the creation of synthetic ER networks based on giant unilamellar vesicles (GUVs), which have a typical size of 10 *µ*m. In contrast to proteo-liposomes, GUVs can be imaged by optical microscopy which allows us to elucidate both the structure and the dynamics of these membrane compartments and to apply the diverse toolbox that has been developed for these vesicles. [30–32]

Using optical microscopy, we demonstrate the importance of the membranes’ lipid composition for the propensity of the GUVs to form ER networks. More precisely, we show that this propensity becomes optimal when the lipid membranes contain a certain mole fraction of cholesterol which has a value between 10 and 15 mol%. This optimal value is comparable to the mole fraction of cholesterol as estimated for ER membranes in vivo. [33] In addition, we demonstrate that synthetic ER networks are obtained from GUV membranes that contain only Sey1 but no Yop1. Our observations imply that the membrane GTPase Sey1 is not only able to induce three-way junctions via membrane fusion but also to generate high membrane curvature, whereas previous studies with proteo-liposomes emphasized that both Sey1 and Yop1 are necessary for the formation of ER networks [22, 23, 25, 28]..

The paper is organized as follows. In the next section, we first look at the effects of GTP on the formation of synthetic ER networks. We show that GUV membranes with atlastin form nanotubes in the absence of GTP but networks with three-way junctions require GTP. In Section 3, we describe the dependence of the network architecture on the mole fraction of cholesterol in the GUV membranes. The propensity for the formation of three-way junctions is observed to attain a maximum for an optimal mole fraction of cholesterol, which has a value between 10 mol% and 15 mol%. In Section 4, we discuss the vesicle morphology when GUV membranes contain the GTPase Sey1 from the yeast *Saccharomyces cerevisiae* instead of the GTPase atlastin from the fruit fly *Drosophila melanogaster*. Surprisingly, we observe the formation of syntheric ER networks for membranes with Sey1 alone even in the absence of Yop1. At the end, we provide a brief summary and outlook as well as a detailed *Methods* section.

## 2 Giant vesicles with membrane-bound atlastin

### 2.1 Binding of atlastin to giant vesicles

The GUVs are prepared from the phospholipid 1-palmitoyl-2-oleoyl-snglycero-3-phosphocholine (POPC) or from binary mixtures of POPC and cholesterol, using the method of electroformation [34, 35]. The GUV membranes are then exposed to membrane proteins such as atlastin in the exterior solution. The molar ratio of lipid to atlastin is 780:1. In order to visualize the binding of the atlastin protein to the GUV membranes, some fraction of the atlastin molecules is labeled by the fluorescent dye Oregon Green (OG), using the labeling protocol described in the *Methods* section.

#### 2.1.1 GUV morphologies in the absence of GTP

When the GUV membranes are exposed to an exterior solution of atlastin in the absence of GTP, the membranes form nanotubes but no three-way junctions. ^†^ Typical GUV morphologies obtained for these experimental conditions are displayed in Figure 1. Panel (a) of this figure shows several GUVs prepared from POPC without cholesterol. The tube diameter is observed to vary between 1.8 and 4 *µ*m. In Figure 1b, the GUV membranes consist of POPC and 10 mol% cholesterol and the tube diameter lies within the range 0.6 - 0.8 *µ*m. This variation of the tube diameter in the same sample is expected to arise from different mole fractions of atlastin as inserted into different GUV membranes.

**Figure 1.**
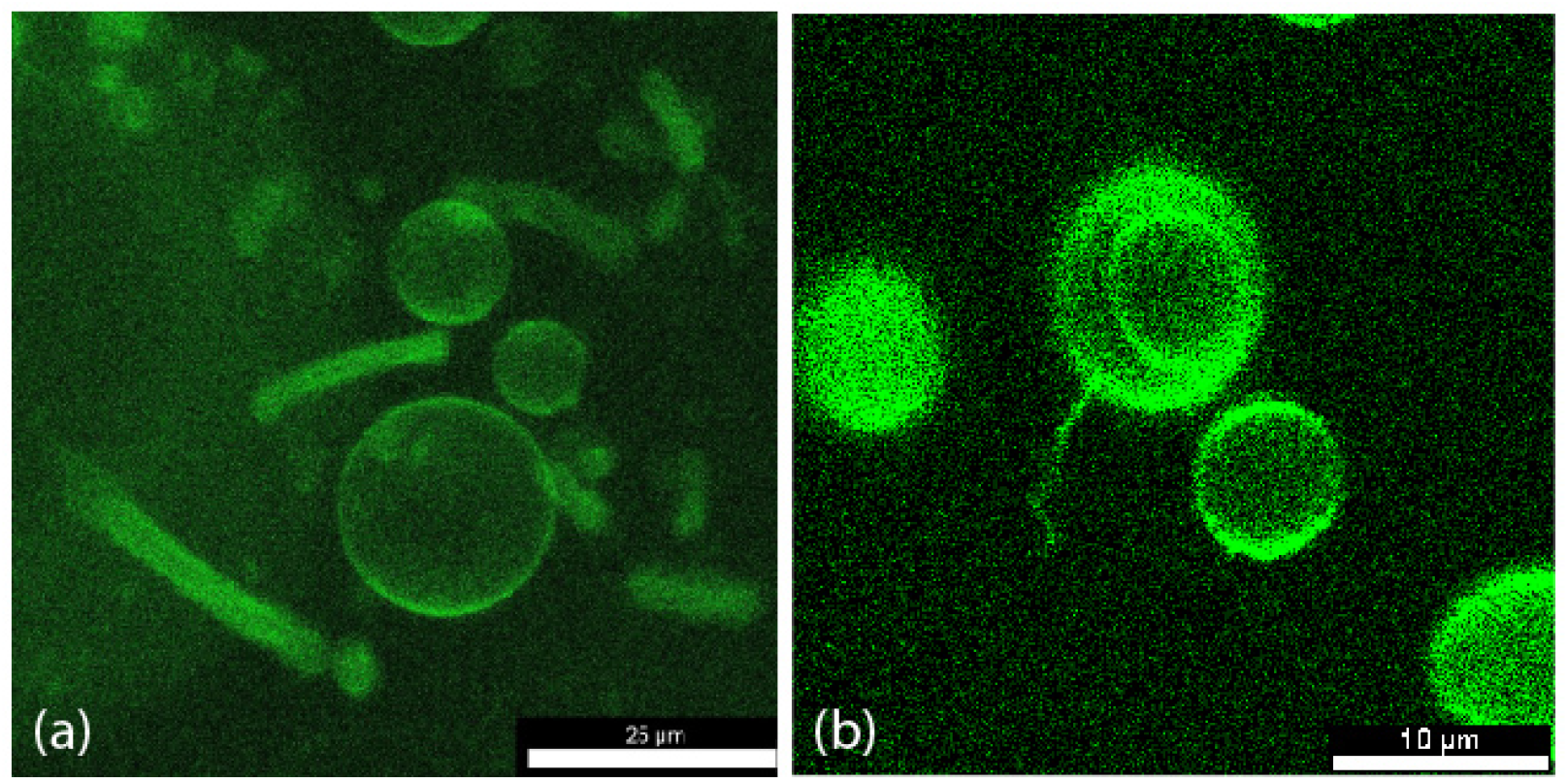
Morphologies of GUV membranes with membrane-bound atlastin in the absence of GTP. The GUVs are exposed to 440 *µ*M atlastin, which is fluorescently labeled by Oregon Green (OG), and 5mM of Mg^2+^ in the exterior solution: (a) Several tubular GUVs with a tube diameter of 1.8 - 4 *µ*m. The lipid bilayers are prepared from the single phospholipid POPC. Scale bar: 25 *µ*m; and (b) A membrane nanotube with a diameter of about 0.7 *µ*m protruding from a GUV with a diameter of about 10 *µ*m. The lipid bilayers are composed of 90 mol% POPC and 10 mol% cholesterol. The tube is very flexible and undergoes strong thermally-excited shape fluctuations, which are more clearly visible in Supplementary Movie 1. Scale bar: 10 *µ*m.

#### 2.1.2 GUV morphology in the presence of GTP

In the presence of GTP, the POPC membranes without cholesterol develop long nanotubes that appear to be rather rigid because they do not exhibit any visible shape fluctuations. One example for this morphology is displayed in Figure 2a. The apparent rigidity of the tubules in the presence of GTP is in strong contrast to the flexible tubes observed in the absence of GTP, see Figure 1b. This apparent rigidity of the nanotubes in Figure 2a provides direct evidence for the presence of a significant membrane tension. Such a tension can arise from the cis-dimerization of two atlastin molecules anchored to the same membrane segment as proposed recently. [19]

**Figure 2.**
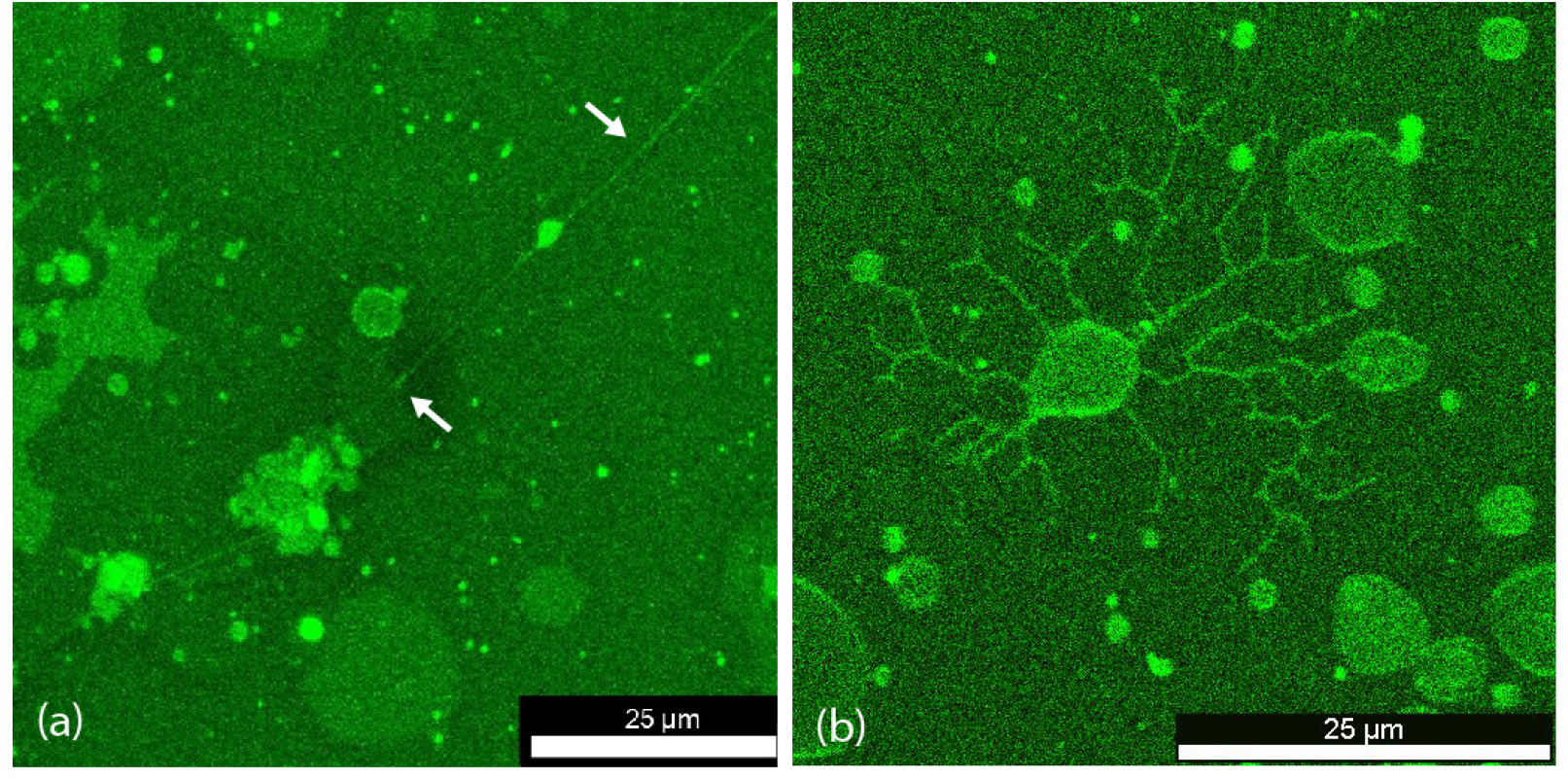
Morphologies of GUV membranes with membrane-bound atlastin in the presence of GTP. In addition to the OG-dyed atlastin, the exterior solution now contains 40 mM GTP and 5mM Mg^2+^ ions: (a) A straight nanotube (white arrows) formed by POPC membranes without cholesterol. An extended view of this nanotube, which has a total length of about 184 *µ*m, is displayed in Supplementary Movie 2; and (b) Nanotubular network formed by a single GUV. The two-component lipid bilayer consists of 90 mol% POPC and 10 mol% cholesterol. The network involves many three-way junctions, each of which connects three nanotubes. A confocal stack of the same field of view is provided by Supplementary Movie 3. Scale bars: 25 *µ*m.

Furthermore, for one-component POPC membranes in the absence of cholesterol, only a small number of three-way junctions have been observed. However, the GUV morphology becomes rather different when we add cholesterol to the POPC membranes. In the latter case, the binary mixture of POPC and cholesterol leads to tubular networks with many three-way junctions connecting three nanotubes, see the example in Figure 2b. In fact, the detailed analysis provided in the next section leads to the conclusion that the occurrence and frequency of three-way junctions varies in a non-monotonic fashion with the mole fraction of cholesterol in the GUV membranes.

Therefore, our study provides strong evidence that the lipid composition is an important control parameter for the formation of synthetic ER networks, in contrast to previous conclusions based on proteo-liposomes for which such a dependence on lipid composition has not been reported. [23] As explained in the *Methods* section, the images of the GUVs in Figures 1 and 2 as well as all other images of GUVs doped with OG-dyed membrane proteins as shown further below have been obtained by confocal microscopy using stroboscopic or pulsed laser light to avoid bleaching of OG, see Figure 7.

## 3 Dependence of synthetic ER networks on cholesterol

To elucidate the dependence of the synthetic ER networks on the mole fraction of cholesterol in a systematic manner, we studied and compared GUV membranes with a mole fraction of 5 mol%, 10 mol%, 15 mol%, and 20 mol% cholesterol. When these GUVs were exposed to an exterior solution of 440 *µ*M atlastin, we observed the formation of reticular networks as displayed in Figure 3, which consists of four panels corresponding to the different mole fractions of cholesterol. Inspection of these different panels reveals that the number of three-way junctions varies strongly with the mole fraction of cholesterol. Indeed, Figure 3a for GUV membranes with 5 mol% exhibits 16 three-way junctions whereas Figure 3b and Figure 3c for 10 and 15 mol% display 76 and 88 three-way junctions, respectively. In addition, a further increase of cholesterol’s mole fraction to 20 mol% as in Figure 3d leads back to the smaller number of 35 three-way junctions. Therefore, the four panels in Figure 3 already indicate that the number of three-way junctions depends on the mole fraction of cholesterol in a non-monotonic manner.

**Figure 3.**
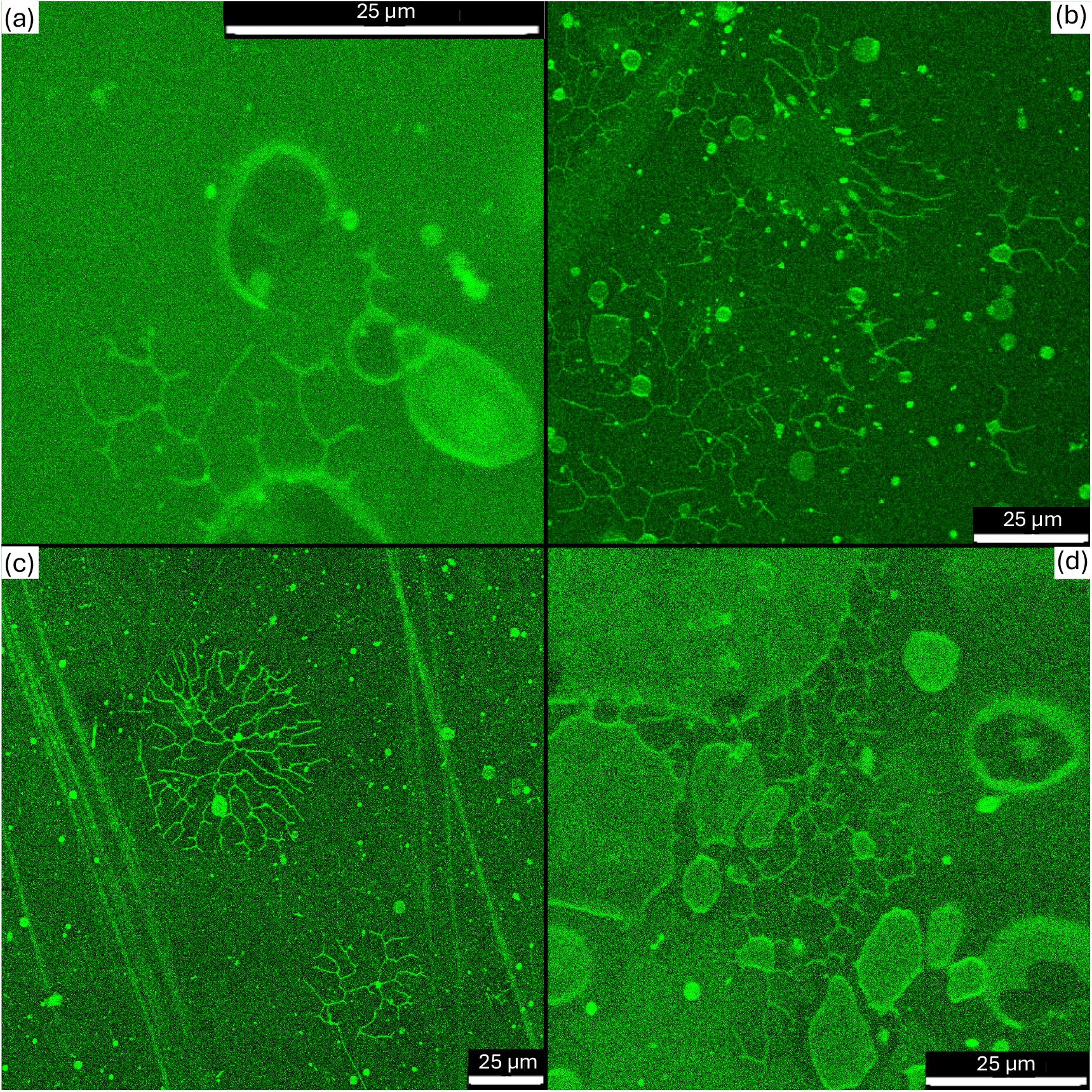
Tubular networks with three-way junctions as observed for POPC membranes with different mole fractions of cholesterol. As in Figure 2, the exterior solution contains 40 mM GTP and 5mM Mg^2+^ in addition to the OG-dyed atlastin: (a) For 5 mol% cholesterol, the GUV form relatively small tubular networks with a few three-way junctions; (b,c) Much more extended networks with many three-way junctions are observed for 10 mol% and 15 mol% cholesterol as in (b) and (c), respectively; and (d) When the mole fraction of cholesterol is increased to 20 mole%, both the size of the networks and the numbers of three-way junctions decrease again. Thus, the propensity to form tubular networks reaches a maximum when the mole fraction of cholesterol is between 10 and 15 mole%. This conclusion is corroborated by the statistics of three-way junctions as displayed in Figure 4.

The qualitative insights arising from Figure 3 are corroborated by the statistics of three-way junctions as depicted in Figure 4 in the form of five bar charts. Each bar chart is composed of four bars in different colors, corresponding to four different mole fractions of cholesterol as defined in the inset of Figure 4. We analyzed each batch of GUVs as obtained for a certain mole fraction of cholesterol by selecting those GUVs that formed membrane nanotubes with one or several three-way junctions and included these GUVs in the data set. The total number of GUVs selected in this way defines the “# of GUVs”, the numerical values of which are provided at the left *y*-axis of Figure 4. The associated bar chart is marked by *†* and denoted by “count”. For each individual GUV from the data set, we determined the number of three-way junctions formed by the GUV. The data set of selected GUVs was then analyzed using four statistical measures as provided by the average number of junctions (Avg) per GUV, the medium (Medium) of the data set, and the standard deviation (Stdev) from the average value. In addition, the bar chart denoted by “Range” describes the difference between the largest and the smallest number of three-way junctions as observed for the individual GUVs in each data set. The numerical values for these statistical measures are displayed at the right *y*-axis of Figure 4.

**Figure 4.**
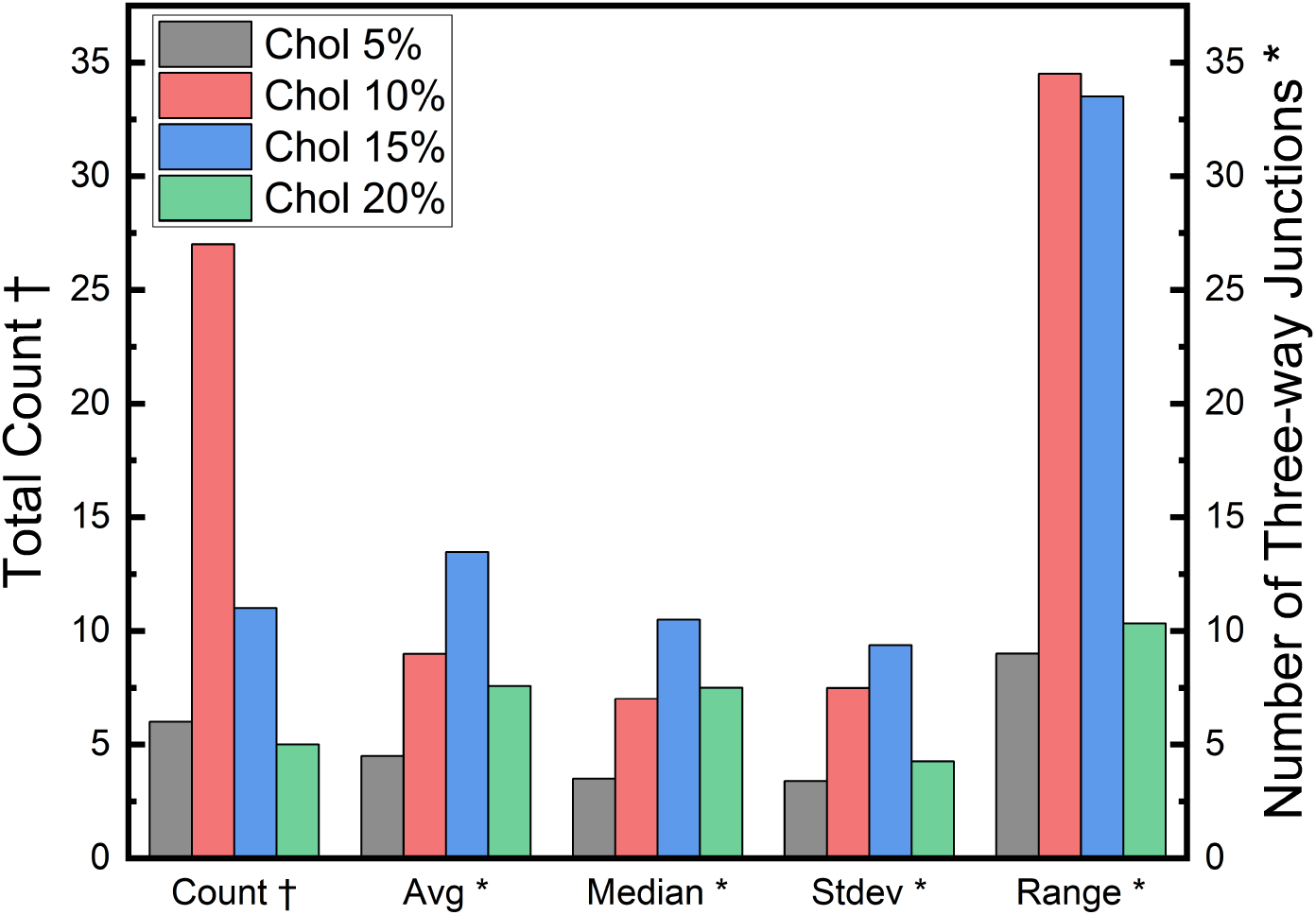
Statistics for the number of three-way junctions as observed for different mole fractions of cholesterol. The statistics is provided in terms of five different bar charts. For each of these charts, the gray, red, green, and blue bars correspond to 5 mole%, 10 mole%, 15 mole%, and 20 mole% cholesterol, respectively. No data are included for 0 mol% cholesterol because, in this case, the number of observed three-way junctions is too small to obtain a meaningful statistics. Count, which is labeled by *†*, refers to the number of GUVs with reticular networks that are included in the data set; the corresponding numerical values are displayed at the left *y*-axis. These datasets are then analyzed in terms of four statistical measures as displayed along the *x*-axis and labeled by *⋆*: Avg stands for the average number of three-way junctions per GUV; Median is the middle value of the data set; and Stdev represents the standard deviation from the average value. Range is the difference between the largest and the smallest number of three-way junctions as observed for the individual GUVs in each data set. The numerical values of these four statistical measures are provided at the right *y*-axis.

The combination of Figures 3 and 4 provides clear evidence that the propensity to form synthetic ER networks varies non-monotonically with the mole fraction of cholesterol and that the frequency of three-way junctions has a maximum for mole fractions between 10 mole% and 15 mole% cholesterol. This optimal value for the synthetic ER networks studied here is comparable to the mole fraction of cholesterol as estimated for ER membranes in vivo. [33]

## 4 Giant vesicles with membrane-bound Sey1

In previous studies with proteo-liposomes, synthetic ER networks have also been obtained for two yeast proteins, the membrane GTPase Sey1 and the membrane protein Yop1 from yeast. [22, 23, 25, 28] In fact, it was thought that Yop1 and Sey1 are both required to obtain such networks, generating a high membrane curvature by Yop1 and forming three-way junctions via membrane fusion by Sey1. In contrast, our studies of GUV membranes, which contain only Sey1, demonstrate that Yop1 is not needed for the formation of synthetic ER networks. One example for such a network formed in the absence of Yop1 is shown in Figure 5a. In this case, the GUV membranes consisted of POPC, 10 mol% cholesterol, and Sey1, which was integrated into the membranes after exposing the vesicles to an exterior aqueous solution of 3.25 *µ*M Sey1.

**Figure 5.**
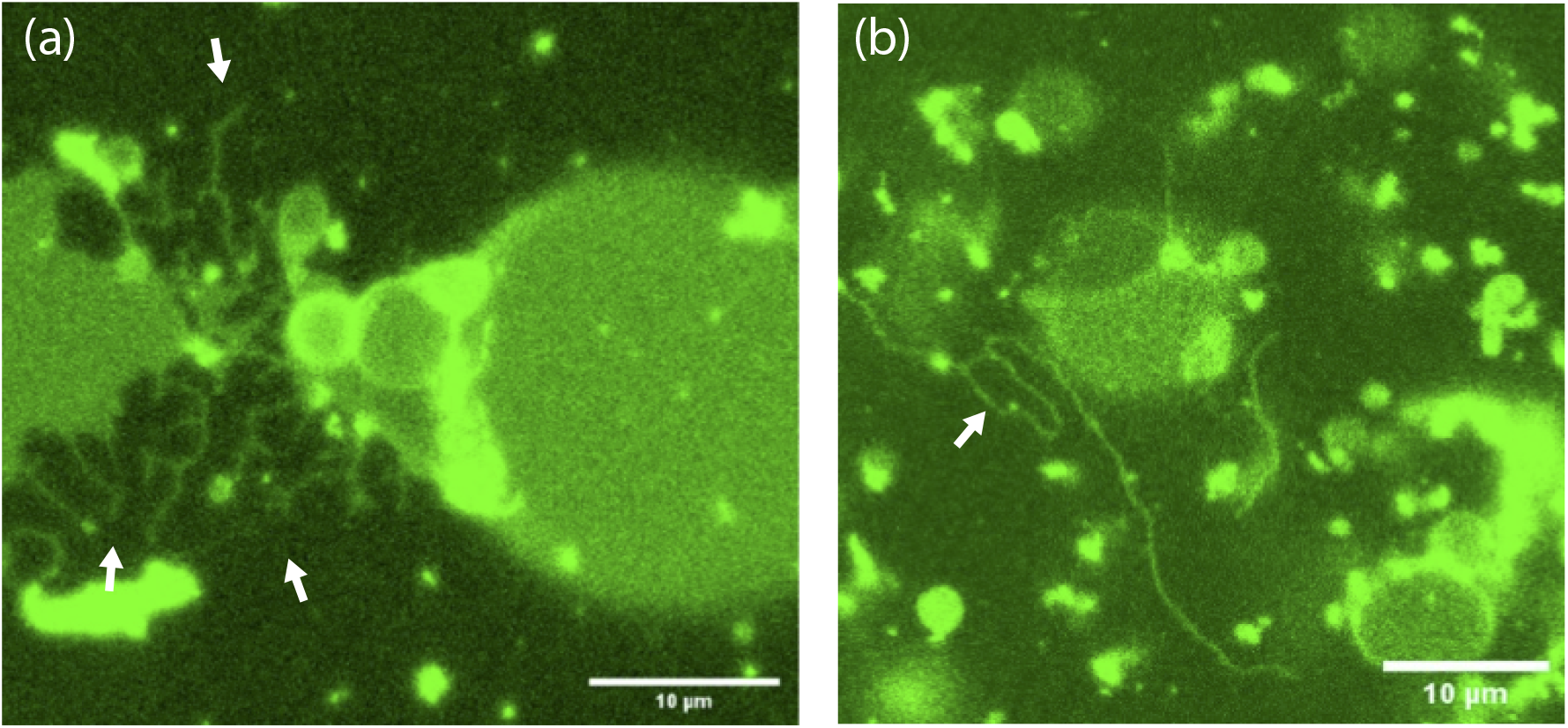
Synthetic ER networks formed by GUVs with the membrane GTPase Sey1 from yeast. The lipid bilayers of the GUVs are prepared from 90 mol% POPC and 10 mol% cholesterol. In addition to Sey1, the exterior solution contains 5 mM of Mg^2+^ ions and 100 mM of GTP: (a) Synthetic ER Networks (bounded by white arrows) are formed when Sey1 is added to the exterior solution at a concentration of 3.25 *µ*M in the absence of the yeast protein Yop1; and (b) The GUVs form long and flexible nanotubes when they are exposed to 3.25 *µ*M Sey1 as well as 1.12 *µ*M Yop1 in the exterior solution. In (b), one of the nanotubes forms a closed loop (white arrow). Both Sey1 and Yop1 are fluorescently labeled by OG, using the same procedure as for atlastin. A larger field of view of the same sample as in (b) is displayed in Suppementary Movie 4.

The synthetic ER networks as displayed in Figure 5a directly demonstrate that the membrane GTPase Sey1 is not only able to create three-way junctions via membrane fusion but also to generate high membrane curvature. On the other hand, the addition of Yop1 to GUV membranes with Sey1 leads to a new tube morphology as shown in Figure 5b. As indicated by the white arrow in this figure, the tubes form loops, which are reminiscent of the irregular polygons as observed for ER networks in vivo. [2, 7] A larger field of view of the same sample as in Figure 5b is displayed in Supplementary Movie 4, which exhibits several loops of membrane nanotubes.

## 5 Summary and outlook

In this paper, we introduce and explore a giant vesicle platform for synthetic endoplasmic reticulum networks. The vesicle membranes contain either pure POPC bilayers or binary lipid mixtures of POPC and cholesterol. When the giant vesicles are exposed to the membrane GTPase atlastin from Drosophila in the exterior solution, the GUVs attain tubular shapes and the GUV membranes form membrane nanotubes (Figure 1) but do not form three-way junctions in the absence of GTP. When GTP is added to the exterior solution, we observe long and rigid nanotubes for pure POPC membranes with atlastin (Figure 2a). Furthermore, for binary lipid mixtures of POPC and cholesterol, the presence of GTP leads to the formation of many three-way junctions connecting three nanotubes (Figure 2b and Figure 3). A systematic comparison of the GUV morphologies as formed by lipid bilayers with 5 mol%, 10 mol%, 15 mol%, and 20 mol% cholesterol reveals that the propensity for the formation of synthetic ER networks varies non-monotonically with the mole fraction of cholesterol (Figure 4) and exhibits a maximum for an optimal mole fraction between 10 and 15 mol%. This optimal value is comparable to the mole fraction of cholesterol as estimated for ER membranes in vivo.

When the GUVs are exposed to an exterior solution with the GTPase Sey1 from yeast, we again observe synthetic ER networks with many three-way junctions (Figure 5a). In contrast, previous synthetic ER networks obtained from proteo-liposomes were based on the insertion of Sey1 and another yeast protein, Yop1, which was believed to be important for the generation of high membrane curvature. Our results directly demonstrate, however, that Yop1 is not necessary for this curvature generation and that Sey1 alone can generate nanotubes with high membrane curvature as well as three-way junctions by membrane fusion.

The giant vesicle platform for synthetic ER networks as described here makes these networks accessible to the versatile toolbox that has been developed for the study of giant vesicles. In fact, many aspects of these nanotubular networks remain to be further elucidated and can be optimized by varying the experimental conditions used for their preparation. For example, it will be rather interesting to use ternary lipid mixtures with cholesterol in order to further optimize the propensity for the formation of these networks.

Another aspect is provided by the network topology. Indeed, most tubular networks discussed here have a tree-like topology without loops. In order to generate more loops, it will be useful to find conditions that increase the local density of the nanotubes. Likewise, another interesting aspect for future studies is how the behavior of the giant vesicle membranes with membrane GTPases such as atlastin or Sey1 changes when one varies the GTP concentration, to which these vesicles are exposed. This concentration is expected, for instance, to affect the undulation spectrum of the membranes because the GTP hydrolysis by atlastin or Sey1 is coupled to the dimerization of the membrane-bound molecules. In particular, one important property of giant vesicle membranes that should be affected by the GTP concentration is the mechanical membrane tension, which can be measured by aspirating the vesicles via micropipettes.

## 6 Methods

### Materials

1-palmitoyl-2-oleoyl-sn-glycero3-phosphocholine (POPC) was from Avanti Polar Lipids (Alabaster, AL). Guanosine triphosphate (GTP) was from Sigma Aldrich. Cholesterol was from Sigma Aldrich. BSA was from Sigma Aldrich. Magnesium chloride was from Sigma Aldrich. Silicone gel was from Carl Roth. Glucose and Sucrose were from Sigma Aldrich. Protein buffer used consisted of 25 mM 4-(2-hydroxyethyl)-1-piperazineethanesulfonic acid, better known as, HEPES pH 7.4, 100 mM KCl, 10% glycerol, 1 mM (Ethylenediaminetetraacetic acid) EDTA, 2 mM *β*-mercaptoethanol. All chemicals were acquired from Thermofisher.

### Protein expression and purification

Atlastin from *Drosophila melanogaster* was expressed using the previously described plasmid pJM681 [21] with an N-terminal GST-tag and a C-terminal His-tag in a pGEX-4T-3 vector. The codon-optimized genes for Sey1 and Yop1 from *Saccharomyces cerevisiae* were ordered from Genscript. Sey1 was expressed using pGEX-6P-3 vector encoding an N-terminal GST-tag followed by a 3C protease site and a C-terminal His-tag. Yop1 was cloned into pOPINF vector [36] coding for an N-terminal His-tag followed by a 3C protease site.

Atlastin was expressed in E. coli BL21-CodonPlus(DE3)-RIL cells in LB medium similar to the previously described protocol [24] but with some modifications. Briefly, the expression was started at an OD600 of about 0.8 with 0.2 mM IPTG for 16-18 h at 18 °C. The cells were harvested, resuspended in buffer A (25 mM HEPES-KOH pH 7.4, 200 mM KCl, 10% (v/v) glycerol, 2 mM DTE) containing 4% Triton X-100, Complete EDTA-free protease inhibitors (Roche), 1 mg/ml lysozyme and 10 µg/ml DNaseI, lysed using a microfluidizer (Microfluidics) and centrifuged at 35,000 rpm for 30 min at 4°C using a 45 Ti rotor (Beckman Coulter). The supernatant was loaded onto a HisTrap FF column (Cytiva), the column was washed with buffer B (25 mM HEPES-KOH pH 7.4, 100 mM KCl, 10% (v/v) glycerol, 2 mM DTE) containing 1% Triton X-100 and 40 mM imidazole and eluted with 500 mM imidazole. The eluted protein was incubated with Glutathione Sepharose 4 Fast Flow resin (Cytiva) for 16-18 h at 4°C. The resin was subsequently washed with buffer B containing 0.1% (or 1%) Triton X-100 and the protein was eluted using the same buffer containing 10 mM reduced glutathione. The fractions containing protein were combined, aliquoted, frozen in liquid nitrogen and stored at -80°C. All protein concentrations were determined using absorbance at 280 nm.

Sey1 was expressed in E. coli Lemo21(DE3) (New England Biolabs) cells in LB medium. Expression was induced at an OD600 of about 0.8 with 0.5 mM IPTG and 1.0 mM L-rhamnose for 16-18 h at 18°C. The cells were harvested by centrifugation and stored at -80°C until further use. The cell pellets were resuspended in buffer A (25 mM HEPES-KOH pH 7.4, 200 mM KCl, 10% (v/v) glycerol, 2 mM DTE) supplemented with Complete EDTA-free protease inhibitors (Roche), 1 mg/ml lysozyme and 20 µg/ml DNaseI and lysed using a microfluidizer (Microfluidics). The cell lysate was then centrifuged at low speed (12,000 g for 10 min at 4°C) to remove cellular debris and unbroken cells. The membranes were sedimented by centrifugation at 45,000 rpm for 45 min at 4°C using a 45 Ti rotor (Beckman Coulter). The pellet was solubilized in buffer A containing 1.5% DDM. The insoluble material was removed by centrifugation at 45,000 rpm for 45 min at 4°C, after which the resulting supernatant was incubated with Ni-NTA resin (Macherey-Nagel) for 16-18 h at 4°C. The resin was washed with buffer B (25 mM HEPES-KOH pH 7.4, 100 mM KCl, 10% (v/v) glycerol, 2 mM DTE) containing 0.1% DDM and 10 mM imidazole. The protein was eluted with buffer B containing 0.1% DDM and 500 mM imidazole and dialyzed against buffer B containing 0.1% DDM. The protein was incubated with Glutathione Sepharose 4 Fast Flow resin (Cytiva) for 1 h at 4°C and the GST-tag was cleaved off (on-column) by incubation with 3C protease for 16-18 h at 4°C. The fractions containing protein were pooled, aliquoted and frozen in liquid nitrogen. Yop1 was expressed in E. coli C43(DE3) cells in TB medium and the induction was performed at an OD600 of about 0.8 with 1.0 mM IPTG for 16-18 h at 16°C.

Yop1 purification was performed similarly to Sey1 with the differences outlined below. After eluting the protein from Ni-NTA resin, 3C protease was added to cleave off the His-tag, and the protein was dialyzed against buffer B containing 0.1% DDM. The sample was loaded onto Ni-NTA column to remove cleaved tag and 3C protease and the flow-through was collected. Finally, the fractions containing protein were combined, aliquoted and frozen in liquid nitrogen.

Protein concentrations were determined using a Denovix DS-11, confocal microscopes used were a Leica TCS SP5 and SP8 and a Zeiss LSM 800. Absorbance measurements were taken on an UltraFast Helios Transient Absorption Spectrometer.

### Labeling of proteins with Oregon Green (OG)

The labeling of atlastin with OG followed the same procedure as used previously. [19] The protein was warmed up to room temperature and added to a sodium bicarbonate solution with a pH above 7.5. OG 488 Carboxylic Acid Succinimidyl Ester from ThermoFisher was added to the protein at a molar ratio of 1:50 or 1:100 and left to sit at room temperature for one hour. After, the protein dye solution was run through a Cytiva PD Minitrap G-25 gravity column, protein buffer was run through the column four times first, then the dyed protein solution was run through the column, collecting into four 2ml tubes, with the second and third tubes being selected for further processing as recommended by the manufacturer per their instructions. Then, using Amicon Ultra Centrifugal Filter, 10 kDa MWCO (molecular weight cut off), the samples were run at 7500 *g* for 10 minutes with the gravitational acceleration *g* = 9.81 m/s^2^ and done multiple times, washing every time with protein buffer until no absorbance value was seen in the supernatant different from the protein buffer being flowed through, as checked by a Denovix DS-11 spectrophotometer. The dyed protein was recollected and suspended with protein buffer. Using a formula from Thermofisher with a correction factor of 0.12 for OG 488, we obtained a labeling degree of about 0.2. However, the measurements of GTP hydrolyis by OG-dyed and unlabeled atlastin as described in the next paragraph indicate a somewhat larger labeling degree.

The procedure for labeling Sey1 and Yop1 with OG 488 Carboxylic Acid Succinimidyl Ester used the same stock as acquired for labeling atlastin and was based on the same procedure as for atlastin.

### GTP hydrolysis by OG-dyed atlastin and unlabeled atlastin

GTP activity was measured using a phosphate assay kit from EnzChek, Invitrogen, as described in [28]. This kit takes any protein or enzyme that releases inorganic phosphate and transforms a certain reagent, 2-amino-6-mercato-7-methylpurine ribonucleoside, which leads to a measurable shift of the absorbance. The phosphate is released from the protein into the solution, where it can react with the reagent through the enzyme purine nucleoside phosphorylase. Both reagent and enzyme are included in the kit.

A phosphate standard was determined using the instructions and materials of the kit. In order to determine the effect of the labeling by OG on the hydrolysis activity of atlastin, both unlabeled atlastin as well as mixtures of OG-dyed atlastin with unlabeled atlastin, as obtained from the labeling procedure, were studied and compared with each other, see Figure 6. The recommended ratios as given in the instructions from EnzChek were followed with a 150 *µ*L cuvette and 166 pmoles of atlastin, thereby preparing 1.11 *µ*M solutions of atlastin. To each solution, 1 *µ*L of 60mM GTP and 5 *µ*L of 5 mM MgCl_2_ were added last and absorbance was measured on an UltraFast Helios Transient Absorption Spectrometer. The adsorbance between 300 and 400 nm was scanned to detect the shift from 330 to 360 nm.

**Figure 6.**
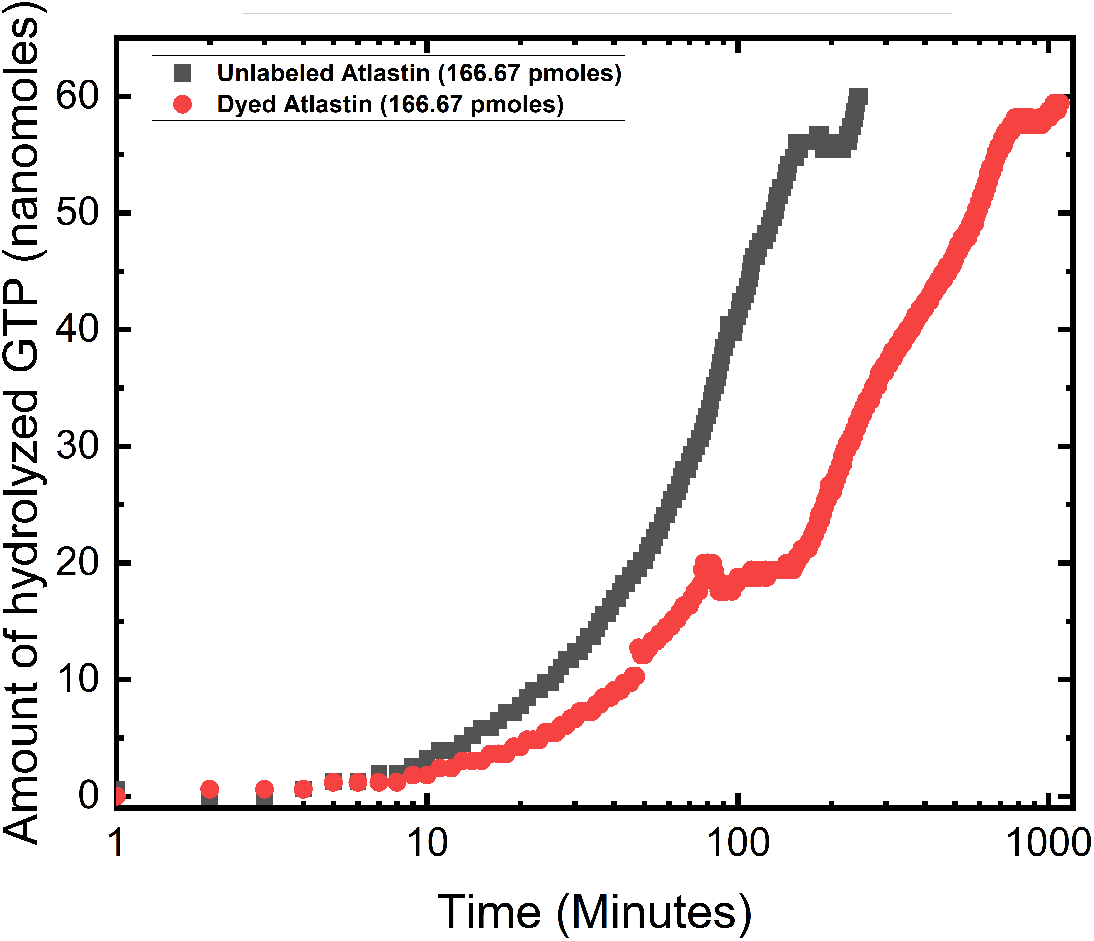
Total amount of GTP hydrolyzed by unlabeled atlastin (black squares) and by mixtures of OG-dyed and unlabeled atlastin (red circles). This measurement was performed over 1000 minutes or 16.7 hours by using a phosphate assay kit as described in the text. The GTP hydrolysis is slowed down by the labeling but the mixture of OG-dyed and unlabeled atlastin eventually hydrolyzes the same total amount of GTP as the solution consisting of unlabeled atlastin only. Time is displayed on a logarithmic scale.

During the first 30 minutes, the 1.11 *µ*M solutions of unlabeled and of mixtures consisting of OG-dyed and unlabeled atlastin hydrolyze 12.39 nmoles and 6.66 nmoles of GTP, respectively, as follows from Figure 6. After 180 minutes, these two types of solutions have hydrolyzed 56.01 nmoles and 23.03 nmoles of GTP. Thus, during the first 30 minutes, the mixtures of OG-dyed and unlabeled atlastin hydrolyze only about half as much GTP as the solutions consisting of only unlabeled atlastin. These data indicate that the labeling degree is about 0.5 if the OG-dyed atlastin lost its ability to hydrolyze GTP. On the other hand, if the OG-dyed atlastin molecules are only partially compromised and still contribute to the GTP hydrolysis, the labeling degree would exceed 0.5.

### Electroformation of GUVs

The GUVs were prepared using the method of electroformation. [34, 35] This method involves using two indium-tin-oxide (ITO)-coated glass plates (PGO GmbH, Iserlohn, Germany) with a piece of conductive copper tape wrapped around the top portion of each plate. Both plates were cleaned with chloroform, then ethanol, and finally water. In a laminar flow cabinet, 10 mM of POPC was mixed with varying molar ratios of cholesterol (0, 5, 10, 15, 20), and/or with varying molar ratios of Oregon Green™ 488 DHPE (0.1, 1, 10) bought from Thermofisher, which was spread across both ITO glass plates, filling up the middle 30% of each plate. These coated plates were put under a vacuum for 30 minutes to evaporate the chloroform used to dissolve the lipid. After removing from vacuum, the two plates were arranged into a closed chamber by using a Teflon spacer as side walls, with the dried lipids on both plates facing the interior of the chamber. The space between the glass plates and the spacer was filled with a sucrose solution of 300 mOsmol (determined by a Gonotec Osmomat Freezing Point Osmometer) and then clamped shut. Using alligator clips, the clamped ITO glasses were connected to an Agilent 33220A Function Generator, which was set to 1.1 V at 10 Hz and run for one hour. Afterwards, the clamps were removed and the aqueous solution with the GUVs was extracted.

### Sample preparation

Sample chambers were prepared from two glass slides that were first washed with deionized water and soap, subsequently dried, and then coated with BSA at 2mg/ml for 5 minutes. On each glass slide, a BSA solution droplet was placed and the contact line between droplet and slide was marked on each slide. Each droplet covered about 35 % of the slide. After 5 minutes, the excess volume of BSA was gently rinsed off with water. Then, silicone gel from Carl Roth was used to create a small doughnut shape on one of the BSA coated glass slides, which was then brought into contact with the other glass slide. All liquids were added to this newly made sample chamber in the hole of the doughnut, and once all liquids had been added, the other slide coated with BSA was put with the BSA facing down on top and slowly pressed down into the silicon gel donut to make a sealed sample chamber. Unless otherwise noted, all experiments were performed with 20 *µ*L of 300 mOsmol glucose, 20 *µ*L of GUV solution in sucrose, 5 *µ*L of 5 mM magnesium chloride, 5 *µ*L of varying GTP solution (ranging from 40 to 100 mM), and varying protein added, typically around 440 *µ*M of atlastin. This was left to sit for 30 to 45 minutes at room temperature before viewing unless otherwise noted.

### Imaging of GUVs by confocal microscopy

Imaging was done with two confocal microscopes, a Leica TCS SP5 or a LeicaTCS SP8. An argon laser was used for both confocals with laser powers ranging from 5 to 20% depending on the experiment with the majority being at 20%. The laser wavelength was 488 nm, with a laser power of 9 to 18%. The laser light was detected via a HyD detector, set to 501 nm or 569 nm. The confocal images were supplemented by phase contrast images. All images were taken with either a Lecia 40*×*, 0.75 aperture dry, or a Lecia 40*×*, 1.2 aperture using oil immersion, where the latter setup was only used for the

SP8. All images have a resolution of either 512 *×*512, 1024 *×*1024, or 2048 *×*2048 pixels, taken at 400, 200, or 100 Hz.

When exposed to continuous laser light, the OG-labeled atlastin undergoes bleaching. The corresponding data are displayed in Figure 7. In order to suppress this bleaching process, individual confocal images were taken with an acquisition time of 1-2 s, separated by much longer time intervals of about 60 s, during which the vesicles were not exposed to any light. The fluorescence intensities of the GUV membranes are deduced by integrating the radial intensity profiles, for which one example is shown in Figure 8. The intensity arising from the OG-labeled atlastin within the membranes is calibrated by comparison with GUV membranes that contain the lipid dye DHPE-OG and, thus, the same fluorophore as the OG-labeled atlastin as in Figure 9.

**Figure 7.**
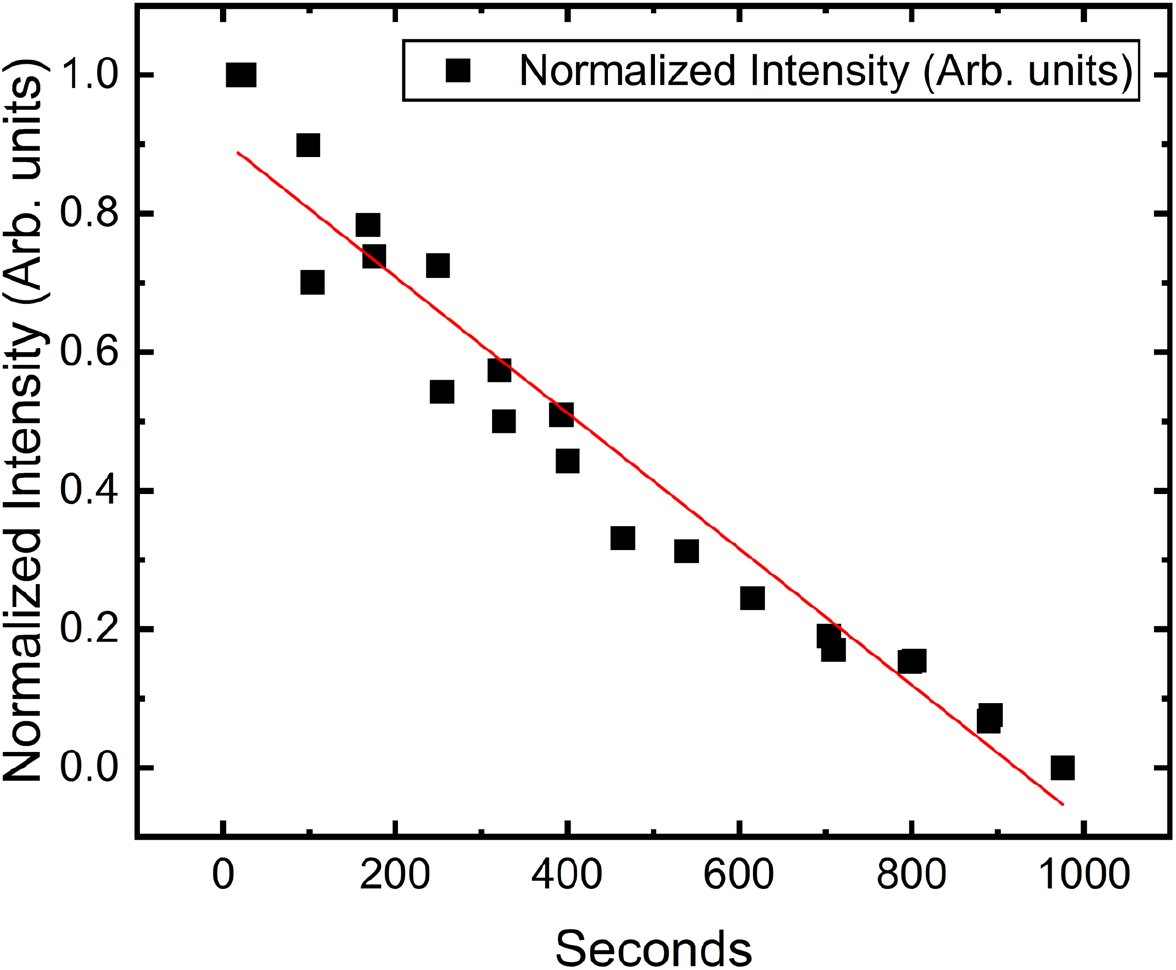
Bleaching of fluorescence intensity of membrane-bound atlastin. **The** GUVs were trapped in a microfluidic chip, loaded with OG-dyed atlastin, and then exposed to constant power of laser light over 1000 seconds. More precisely, the bleaching experiment was done with confocal microscope settings for the image acquisition of 512 *×* 512 pixels at 200 Hz with 9% 488 laser power. In order to strongly reduce these bleaching effects, all other confocal images reported here were obtained with pulsed laser light, that is, acquiring individual images over short time intervals of 1-2 seconds, separated by dark periods of about 60 seconds.

**Figure 8.**
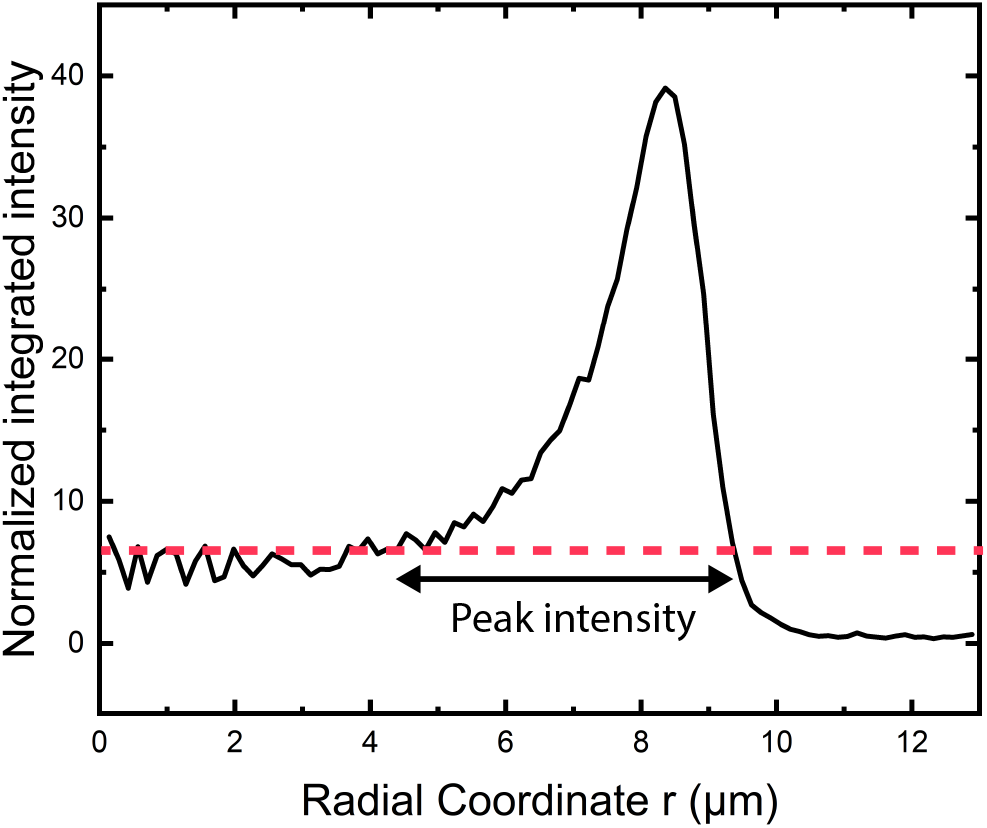
Radial intensity profile of a GUV membrane with 90 mol% POPC and 10 mol% cholesterol. The GUV is exposed to an exterior solution with 440 *µ*M OG-labeled atlastin and 40 mM GTP. The center of the spherical vesicle defines the origin of the radial coordinate, *r* = 0. The broken horizontal line (red) at *y* = 6.25 is used to define the lower and upper bounds for the integration over the peak intensity as indicated by the double arrow.

**Figure 9.**
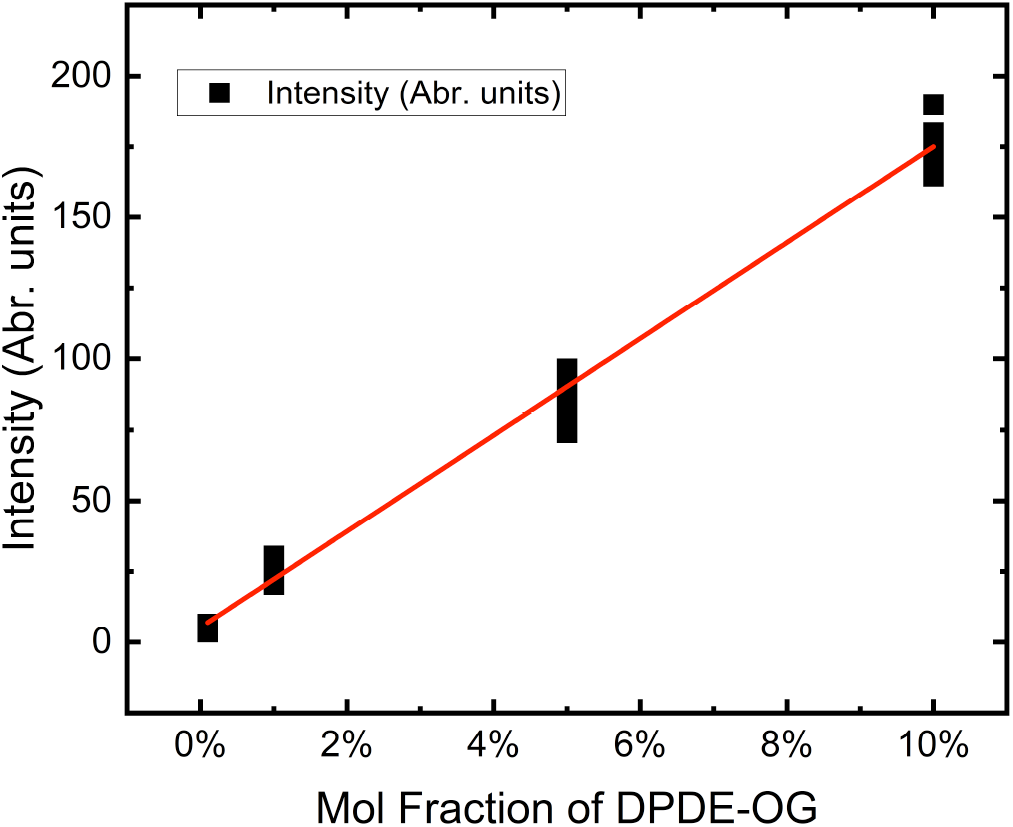
Fluorescence intensity of GUV membranes consisting of POPC, 10 mol% cholesterol, and the fluorescent lipid dye DHPE-OG. The four data points correspond to 0 mol%, mol%, 1 mol%, 5 mol%, and 10 mol% DHPE-OG. The red line is a linear fit obtained via Origin. These data can be used to calibrate the fluorescence intensity of GUV membranes consisting of POPC, 10 mol% cholesterol, and varying amounts of OG-labeled atlastin.

### Bleaching of OG-dyed atlastin

The membrane-bound atlastin labeled by OG undergoes bleaching when exposed to laser light at constant laser power. The bleaching process was studied in a microfluidic chip, which was designed to trap the GUVs as in previous studies [37,38]. The chip was passivated with BSA by adding 5 *µ*L of 2mg/mL BSA solution into the chip and spinning it in a centrifuge at 200 g for 10 minutes.

After passivating with BSA and then washing out with 300mOSmol/kg glucose, GUVs with 90 mol% POPC and 10 mol% cholesterol were flushed into the chamber and confined into rectangular traps formed by PDMS posts of dimensions 40*×* 40 *µ*m^2^, with an average spacing of 2.5 *µ*m between them. GUVs were flushed in for 30 minutes at a rate of 1 *µ*L/min. Afterwards, atlastin was added where 15 ng of atlastin were diluted in 100 *µ*L of 300 mOSmol/kg glucose. This was flushed through the chip for 30 minutes to let the atlastin integrate into the membrane. Afterwards 10 *µ*L of 5mM MgCl2 and 10 *µ*L of 60mM GTP in 100 *µ*L of 300mOSmol/kg glucose was flushed through the system at a rate of 1 *µ*L/min. The GUV solution was exposed to constant laser power and the fluorescence intensities of atlastin were measured for a region of interest that was imaged with a Leica SP8 confocal taking *z*-stacks with 31 frames. Bleaching rate was calculated from these stacks over the course of nearly 1000 s or 17 minutes as in Figure 7. A single region was identified, and using nearby marking, the correct *z*-level was found for each continuous z stack. This bleaching experiment was done with confocal microscope settings for the image acquisition of 512 *×* 512 pixels at 200 Hz with 9 % laser power.

The data in Figure 7 were obtained exposing the atlastin-doped GUVs to constant laser light (continuous wave laser operation). In order to strongly reduce the bleaching of OG-labeled atlastin, all other experimental data were obtained acquiring individual images (one image acquired over 1-2 s) separated by dark periods of about 60 s.

### Fluorescence intensity of GUV membranes

For image processing, the confocal images were loaded into ImageJ with Fiji as TIFF files. The intensities were found for each *z*-stack using Radial Profile Extended Plugin from Github repository [39]. To generate a radial intensity profile, a circular region was created in imageJ. This circular region was placed over the GUV of interest, being about 10 - 20% larger than the GUV, so that the background intensity would show up in the radial intensity profile plot. The circular region of interest was located in such a way that its center was located at the center of the GUV. The membrane intensity was then obtained by integrating the peak of the intensity profile between a lower and upper bound as described previously. [40] One example for such a radial intensity profile and the associated lower and upper bound for the integration of the peak intensity are displayed in Figure 8.

### Calibration of fluorescence intensity

To obtain a reference value for the fluorescence intensity of OG within a lipid bilayer, we studied GUV membranes with the lipid dye Oregon Green 488 1,2-Dihexadecanoyl-sn-Glycerin-3-Phosphoethanolamin (DHPE-OG) from Thermofisher in the absence of OG-labeled atlastin. We determined the fluorescence intensity for several mole fractions of DHPE-OG as shown in Figure 9. These data can be used to calibrate the fluorescence intensity of GUV membranes with the lipid dye DHPE-OG replaced by OG-dyed atlastin bound to the membrane.

## Acknowledgments

This work was supported by the Max Planck Institute of Colloids and Interfaces as well as by the Max Planck Society and the Dieter Schwarz foundation via the Max Planck School “Matter to Life”. Open access funding is provided by the Max Planck Society.

## Author contributions

R.L. designed the project. C. V. performed the experiments. S. P. developed the initial experimental protocols. T.-W. S. did control experiments. M. T. expressed and purified the membrane proteins. J. S. and R. D. provided experimental advice R. L. and C. V. wrote the manuscript. All authors discussed the results and commented upon the manuscript.

## Data availability statement

All data that support the findings of this study are included within the article (and any supplementary files).

## Supplementary informations

The online version contains four Supplementary Movies 1 - 4, available at https://doi.org/…

## Author information

## Footnotes

† The same behavior is observed for exterior solutions without magnesium.

## References

[1] B. Alberts, D. Bray, J. Lewis, M. Raff, K. Roberts, and J. Watson: Molecular biology of the cell (Garland Publ., New York, NY, USA), third edition (1994).

[2] C. J. Obara, A. S. Moore, and J. Lippincott-Schwartz: Structural Diversity within the Endoplasmic Reticulum - From the Microscale to the Nanoscale. Cold Spring Harb. Perspect. Biol. 15, a041259 (2023).

[3] R. E. Pagano, K. J. Longmuir, O. C. Martin, and D. K. Struck: Metabolism and Intracellular Localization of a Fluorescently Labeled Intermediate in Lipid Biosynthesis within Cultured Fibroblasts. J. Cell Biol. 91, 872–877 (1981).

[4] M. Terasaki, J. Song, J. F. Wong, M. J. Weiss, and L. B. Chen: Localization of Endoplasmic Reticulum in Living and Glutaraldehyde-Fixed Cells with Fluorescent Dyes. Cell 38, 101–108 (1984).

[5] C. Lee and L. B. Chen: Dynamic Behavior of Endoplasmic Reticulum in Living Cells. Cell 54, 37–46 (1988).

[6] O. Baumann and B. Walz: Endoplasmic Reticulum of Animal Cells and Its Organization into Structural and Functional Domains. Int. Rev. Cyt. 205, 149–214 (2001).

[7] A. M. Valm, S. Cohen, W. R. Legant, J. Melunis, U. Hershberg, E. Wait, A. R. Cohen, M. W. Davidson, E. Bethig, and J. Lippincott-Schwartz: Applying systems-level spectral imaging and analysis to reveal the organelle interactome. Nature 546, 162–167 (2017).

[8] M. Puhka, H. Vihinen, M. Joensuu, and E. Jokitalo: Endoplasmic reticulum remains continuous and undergoes sheet-to-tubule transformation during cell division in mammalian cells. J. Cell Biol. 179, 895–909 (2007).

[9] L. Lu, M. S. Ladinsky, and T. Kirchhausen: Cisternal Organization of the Endoplasmic Reticulum during Mitosis. Mol. Biol. Cell 20, 3471–3480 (2009).

[10] M. Puhka, M. Joensuu, H. Vihinen, I. Belevich, and E. Jokitalo: Progressive sheet-to-tubule transformation is a general mechanism for endoplasmic reticulum partitioning in dividing mammalian cells. Mol. Biol. Cell 23, 2424–2432 (2012).

[11] S. Kors and A.-L. Schlaitz: Dynamic remodelling of the endoplasmic reticulum for mitosis. J. Cell Science 137, jcs261444 (2024).

[12] M. Terasaki: Axonal endoplasmic reticulum is very narrow. J. Cell Sci. 131, jcs210450 (2018).

[13] G. K. Voeltz, W. A. Prinz, Y. Shibata, J. M. Rist, and T. A. Rapoport: A Class of Membrane Proteins Shaping the Tubular Endoplasmic Reticulum. Cell 124, 573–586 (2006).

[14] J. Hu, Y. Shibata, C. Voss, T. Shemesh, Z. Li, M. Coughlin, M. M. Kozlov, T. A. Rapoport, and W. A. Prinz: Membrane Proteins of the Endoplasmic Reticulum Induce High-Curvature Tubules. Science 319, 1247–1250 (2008).

[15] R. Lipowsky: Spontaneous tubulation of membranes and vesicles reveals membrane tension generated by spontaneous curvature. Faraday Discuss. 161, 305–331 (2013).

[16] Y. Li, R. Lipowsky, and R. Dimova: Membrane nanotubes induced by aqueous phase separation and stabilized by spontaneous curvature. Proc. Nat. Acad, Sci. USA 108, 4731–4736 (2011).

[17] M. J. Dayel, E. F. Y. Hom, and A. S. Verkman: Diffusion of Green Fluorescent Protein in the Aqueous-Phase Lumen of Endoplasmic Reticulum. Biophys. J. 76, 2843–2851 (1999).

[18] D. Holcman, P. Parutto, J. E. Chambers, M. Fantham, L. J. Young, S. J. Marciniak, C. F. Kaminski, D. Ron, and E. Avezov: Single particle trajectories reveal active endoplasmic reticulum luminal flow. Nat. Cell Biol. 20, 1118–1125 (2018).

[19] R. Lipowsky, S. Pramanik, A. S. Benk, M. Tarnawski, J. P. Spatz, and R. Dimova: Elucidating the Morphology of the Endoplasmic Reticulum: Puzzles and Perspectives. ACS Nano 17, 11957–11968 (2023).

[20] L. Dreier and T. A. Rapoport: In Vitro Formation of the Endoplasmic Reticulum Occurs Independently of Microtubules by a Controlled Fusion Reaction. J. Cell Biol. 148, 883–898 (2000).

[21] G. Orso, D. Pendin, S. Liu, J. Tosetto, T. J. Moss, J. E. Faust, M. Micaroni, A. Egorova, A. Martinuzzi, J. A. McNew, and A. Daga: Homotypic fusion of ER membranes requires the dynamin-like GTPase Atlastin. Nature 460, 978–983 (2009).

[22] J. Hu, Y. Shibata, P.-P. Zhu, C. Voss, N. Rismanchi, W. A. Prinz, T. A. Rapoport, and C. Blackstone: A Class of Dynamin-like GTPases Involved in the Generation of the Tubular ER Network. Cell 138, 549–561 (2009).

[23] R. E. Powers, S. Wang, T. Y. Liu, and T. A. Rapoport: Reconstruction of the tubular endoplasmic reticulum network with purified components. Nature 543, 257–272 (2017).

[24] M. A. Betancourt-Solis, T. Desai, and J. A. McNew: The atlastin membrane anchor forms an intramembrane hairpin that does not span the phospholipid bilayer. J. Biol. Chem. 293, 18514–18524 (2018).

[25] N. Wang and T. A. Rapoport: Reconstituting the reticular ER network - mechanistic implications and open questions. J. Cell Sci. 132, jcs227611 (2019).

[26] T. Y. Liu, X. Bian, S. Sun, X. Hu, R. W. Klemm, W. A. Prinz, T. A. Rapoport, and J. Hu: Lipid interaction of the C terminus and association of the transmembrane segments facilitate atlastin-mediated homotypic endoplasmic reticulum fusion. Proc. Natl. Acad. Sci. USA 109, E2146–E2154 (2012).

[27] T. Y. Liu, X. Bian, F. B. Romano, T. Shemesh, T. A. Rapoport, and J. Hu: Cis and trans interactions between atlastin molecules during membrane fusion. Proc. Natl. Acad. Sci. USA pp. E1851–E1860 (2015).

[28] K. Anwar, R. W. Klemm, A. Condon, K. N. Severin, M. Zhang, R. Ghorlando, J. Hu, T. A. Rapoport, and W. A. Prinz: The dynamin-like GTPase Sey1p mediates homotypic ER fusion in S. cerevisiae. J. Cell Biol. 197, 209–217 (2012).

[29] X. Bian, R. W. Klemm, T. Y. Liu, M. Zhang, S. Sun, X. Sui, X. Liu, T. A. Rapoport, and J. Hu: Structures of the atlastin GTPase provide insight into homotypic fusion of endoplasmic reticulum membranes. Proc. Natl. Acad. Sci. USA 108, 3976–3981 (2011).

[30] R. Lipowsky and E. Sackmann (eds.): Structure and Dynamics of Membranes, volume 1 of Handbook of Biological Physics (Elsevier, Amsterdam) (1995).

[31] P. Bassereau and P. Sens (eds.): Physics of Biological Membranes (Springer Nature Switzerland AG, Cham, Switzerland) (2018).

[32] R. Dimova and C. Marques (eds.): The Giant Vesicle Book (CRC Press, Boca Raton, FL, USA) (2019).

[33] G. van Meer, D. R. Voelker, and G. W. Feigenson: Membrane lipids: where they are and how they behave. Nature Rev.: Mol. Cell Biol. 9, 112–124 (2008).

[34] M. Angelova and D. Dimitrov: Liposome electroformation. Faraday Discuss. Chem. Soc.81, 303–311 (1986).

[35] R. Dimova, P. Stano, C. M. Marques, and P. Walde: Preparation methods for giant unilamellar vesicles. In R. Dimova and C. M. Marques (eds.), The Giant Vesicle Book, chapter 1 (CRC Press) (2019).

[36] N. S. Berrow, D. Alderton, S. Sainsbury, J. Nettleship, R. Assenberg, N. Rahman, D. I. Stuart, and R. Owens: A versatile ligation-independent cloning method suitable for high-throughput expression screening applications. Nucleic Acids Res. 35, e45 (2007).

[37] T. Robinson, P. Kuhn, K. Eyer, and P. S. Dittrich: Microfluidic trapping of giant unilamellar vesicles to study transport through a membrane pore. Biomicrofluidics 7, 044105 (2013).

[38] N. Yandrapalli and T. Robinson: Ultra-high capacity microfluidic trapping of giant vesicles for high-throughput membrane studies. Lab Chip 19, 626–633 (2019).

[39] P. Carl: Radial Profile Extended (2023). Webpage of GitHub repository at https://github.com/phcarl/Radial-Profile-Extended

[40] S. Pramanik, J. Steinkühler, R. Dimova, J. P. Spatz, and R. Lipowsky: Binding of His-tagged fluorophores to lipid bilayers of giant vesicles. Soft Matter 18, 6372–6383 (2022).

